# Neural population variance explains adaptation differences during learning

**DOI:** 10.1101/2024.10.01.616115

**Authors:** Hannah M. Stealey, Yi Zhao, Hung-Yun Lu, Enrique Contreras-Hernandez, Yin-Jui Chang, Philippe Tobler, Jarrod A. Lewis-Peacock, Samantha R. Santacruz

## Abstract

Variability, a ubiquitous feature of neural activity, plays an integral role in behavior. However, establishing a causal relationship between neural signals and behavior is difficult. By defining a mathematical mapping between neural spiking activity and behavior, we investigate the role of spiking variability in adaptation during a brain-computer interface (BCI) behavior in male rhesus macaques (*Macaca mulatta,* n=2). Recent BCI evidence demonstrates that creating novel neural patterns is harder than repurposing existing patterns to respond to changes in external input. However, what limits the ability to repurpose, or adapt, patterns under different magnitudes of change is less well-characterized. Here, we present evidence that variance in neural spiking activity reveals differences in learnability between easy and hard adaptation conditions and across sessions. Furthermore, our study illuminates the limitations in neural changes underlying behavior within a neurofeedback paradigm.

**Significance Statement:** Variability in neural activity is a major driver of behavioral variability, though it is unclear how variability is balanced with stable neural activity as new behaviors become more practiced. By using a brain-computer interface methodology, we define a mathematical mapping between neural spiking activity and a behavioral control signal. Through thoughtful manipulation of this mapping, we incite the subjects (rhesus macaques) to learn and adapt neural activity to regain behavioral proficiency. We find that metrics of neural population variability are differentially modulated depending on difficulty of the imposed manipulation. Our exciting results provide important implications for brain-computer interface applications as well as our understanding of learning and adaptation more broadly. Our work represents an important step forwards towards understanding population neural dynamics in this critical component of behavior.

## Introduction

Regardless of neural scale (e.g., single neurons, populations of neurons, neural systems), variability is a common feature of neural signals (Faisal et al., 2008; Garrett et al., 2018; Uddin, 2020; Waschke et al., 2021). Neural variability refers to the inherent variability in the activity of neurons, even when they are exposed to the same external input or behavior. Variability in neural activity is a major driver of behavioral variability. While it might seem counterintuitive, behavioral variability is thought to be important when learning new behaviors (Wu et al., 2014; Dhawale et al., 2017). Once a skill is acquired and stable behavior is desired, however, variability in behavior is often regarded as unwanted “noise” (Harris and Wolpert, 1998; Dhawale et al., 2017; Sternad, 2018). Still, the ability to adapt learned behaviors on demand is crucial for living in an ever-changing world. Altogether, a more nuanced approach of investigating neural variability underlying behavior may be the key to untangling the role variability plays in adaptive learning (Krakauer and Mazzoni, 2011; Dhawale et al., 2017; Krakauer et al., 2019).

Because neural variability can encode a variety of brain states, disentangling the causal role of neural variability in task-related output is challenging. The challenge stems from the difficulty in observing all high-dimensional neural patterns that ultimately produce a single, low-dimensional task-relevant behavior through a variable motor system (Golub et al., 2018; Zippi et al., 2022). Using brain-computer interfaces (BCIs), researchers circumvent this challenge by creating a causal, mathematical mapping (“decoder”) between the activity of small populations of neurons and a behavioral output. By definition, BCIs directly establish the relationship between a specified group of neurons and behavioral variables. Thus, all task-relevant neurons are known, and the activity of these neurons can be observed throughout the BCI behavior.

BCI studies in non-human primates (NHPs) have established key neural features of BCI learning and adaptation. Neural activity patterns span an *n*-dimensional space in which *n* is the number of neurons. However, intrinsic neural activity patterns tend to reside in a lower-dimensional portion of this space (“neural subspace” or “intrinsic manifold”) (Sadtler et al., 2014). During *de novo* BCI learning, generating novel neural patterns of co-activation outside of the neural subspace occurs gradually over days (Ganguly and Carmena, 2009; Oby et al., 2019; Zippi et al., 2022) but not significantly within a single session (Sadtler et al., 2014; Oby et al., 2019). Over the learning process, pairwise correlations between neural decoder units (i.e., neurons) increase (Ganguly and Carmena, 2009). Additionally, patterns of neural population activity exhibit increasing levels of shared variance as a function of learning over the timespan of days (Athalye et al., 2017; Zippi et al., 2022). When adapting learned or naturally occurring patterns within the neural subspace as a result of a perturbation to the decoder mapping, repurposing these intrinsic patterns (neural “reassociation” (Sadtler et al., 2014; Golub et al., 2018)) is easier (i.e., can be accomplished on a shorter timescale) than generating novel patterns that reside outside of the known neural subspace (Sadtler et al., 2014; Golub et al., 2018; Hennig et al., 2018; Oby et al., 2019). While this body of work provides a key insight into what neural patterns decoder populations can readily learn and proposes a neural mechanism for adaptation, the differences in the ability of decoder units to adapt patterns within the known neural subspace are not well understood. To study this gap, neural spiking activity in decoder populations needs to be investigated in a framework that explains the capacity of neural units to adapt existing neural patterns.

To investigate how neural variability contributes to behavioral adaptation, we implemented a BCI perturbation task with NHPs. Specifically, we perturbed the decoder using a visuomotor rotation (VMR) in which the behavioral output (2D cursor movements on a computer screen) appeared rotated by an angle. The magnitude of the angle controlled task difficulty (Jarosiewicz et al., 2008; Chase et al., 2012; Vyas et al., 2018). Importantly, VMR perturbations provide a framework in which a feasible solution is reassociation of existing neural patterns with new directions of movement; this can be learned within a day (Hennig et al., 2018).

## Results

### Experimental setup: center-out visuomotor rotation brain-machine interface task

We trained male Rhesus macaque monkeys (*Macaca mulatta*, n=2) in a center-out, VMR BCI task (Ganguly and Carmena, 2009, 2010; Chase et al., 2012; Sadtler et al., 2014; Athalye et al., 2017; Golub et al., 2018; Hennig et al., 2018; Oby et al., 2019; Zippi et al., 2022). During this task, each subject volitionally modulated spiking activity of neural units from motor cortical areas to control a cursor shown on a computer screen (**Figure 1A**). Neural population spiking activity was mapped to cursor velocity through a linear Kalman filter (KF) decoder, which produced a control signal to update cursor position every 100 milliseconds (10 Hz update rate) (Wu et al., 2006; Dangi et al., 2014). During the task, the subjects received continuous visual feedback of the current cursor position.

**Figure 1.**
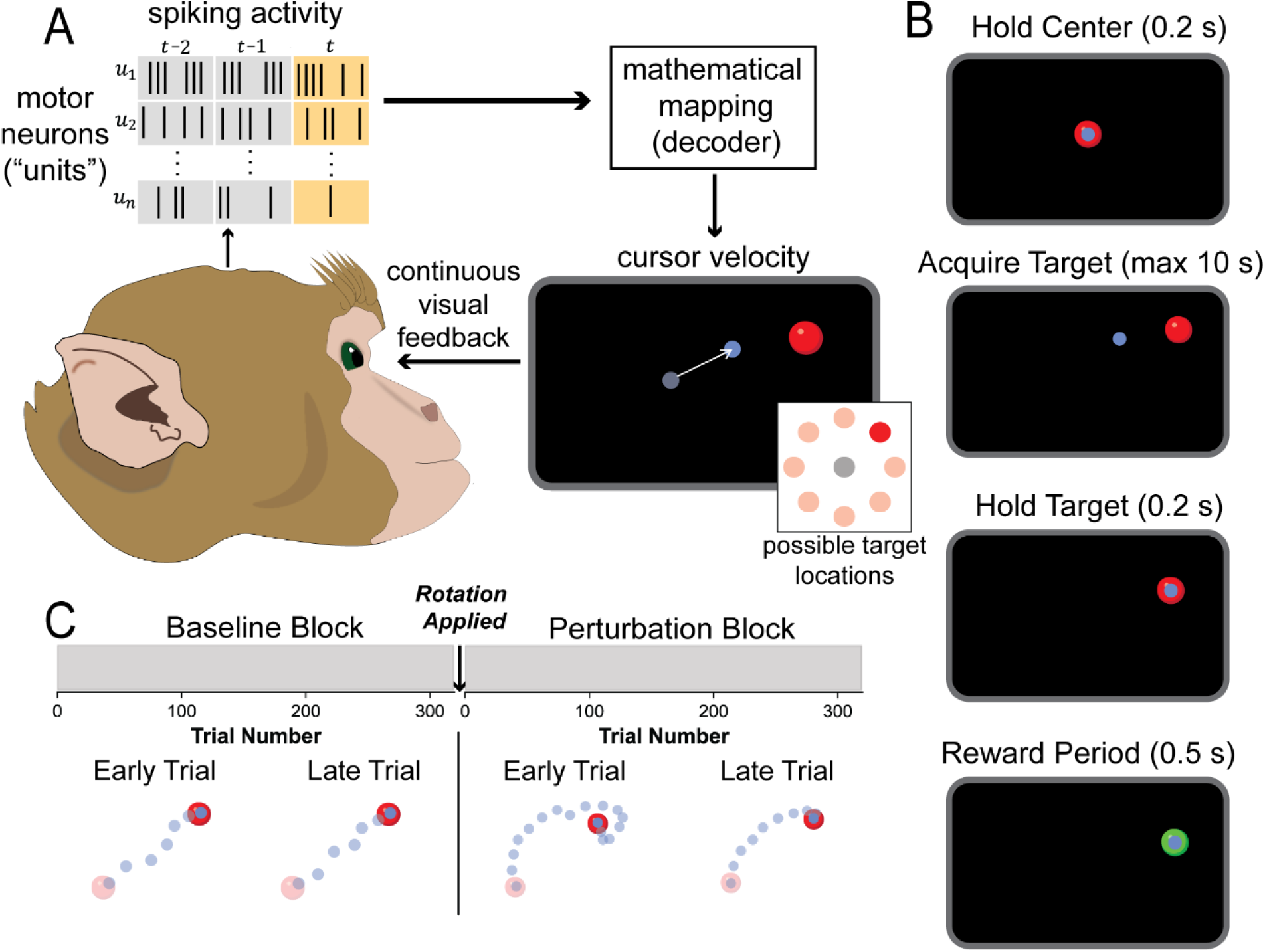
Experimental setup for BCI task with a visuomotor rotation perturbation. (A) Brain-computer interface paradigm. Subjects modulated neural spiking activity from motor cortical (forelimb) areas to control the velocity of a computer cursor in two dimensions. Time bins (t) of neural unit (*u*_*i*_) spiking activity were translated through a linear KF “decoder” to update the cursor position. Visual feedback allowed subjects to learn to volitionally produce neural patterns that control cursor velocity proficiently. (B) Single trial structure. The subject held the cursor (small blue circle) at the center target (large red circle; Hold Center) then acquired the peripheral target within 10 s (Acquire Target). After holding the cursor at the target (Hold Target), the target turned green, and the subject received a small liquid reward to reinforce this behavior (Reward Period). (C) Single session structure (blocks of trials). During the baseline block, subjects operated a decoder that was trained and tuned to their neural activity each day. Notably, early and late baseline trials contained cursor trajectories that lead relatively straight to the peripheral target (dark red circle) in similar amounts of cursor updates (i.e. time). At the onset of the subsequent perturbation block, a rotation perturbation was applied. This impacted the length and linearity of the path the cursor traveled, as well as the time to complete a trial. To regain proficient control, the monkeys intentionally altered their neural spiking activity to counteract (adapt to) the perturbation (Perturbation – Late Trial). Two rotation conditions (Easy/50°, Hard/90°) were investigated over the course of this study, with only one rotation perturbation presented per session.

All trials followed the same sequence of events (**Figure 1B**). At the start of each trial, the subject held the cursor within a circular target located at the center of the computer screen for 200 milliseconds. Once the subject successfully completed the hold, a peripheral target appeared. The subject was allotted 10 seconds to move the cursor to the peripheral target. Following target acquisition, the subject was required to keep the cursor within target bounds for 200 milliseconds. The 200-millisecond hold time for the center and peripheral holds started once at least half of the cursor’s area overlapped with the target. After a successful hold period, the subject received a small liquid reward to reinforce the behavior (∼100 µL of apple juice).

On a single trial, one of eight radially-dispersed targets was presented 10 centimeters from the center target (**Figure 1A**). The subject completed a single trial to each of the eight targets in a pseudorandom order prior to the target locations repeating. For the remainder of this paper, consecutive, non-overlapping groups of eight trials are referred to as a “set of trials”. Additionally, we refer to the order of each trial set as the trial set number (TSN).

In each session, subjects completed a baseline block first followed by a perturbation block. Each block contained 42 sets of trials (i.e., 336 individual trials) for analysis. In the first block (**Figure 1C**), subjects used a decoder that was trained and re-calibrated every behavioral session (Dangi et al., 2014). The purpose of the baseline block was to establish the subject’s daily baseline behavioral performance and neural spiking dynamics. At the start of the perturbation block, we instantiated a VMR perturbation to the decoder (**Figure 1C**). In a given day, the rotation angle remained fixed at the onset of the perturbation block at either 50° or 90°. The 50° perturbation served as an “easy” rotation, whereas the 90° perturbation served as a “hard” rotation. These rotation conditions were selected based on VMR literature in NHPs (Jarosiewicz et al., 2008; Chase et al., 2012; Vyas et al., 2018) and humans (Krakauer et al., 2000; Tanaka et al., 2009; Huang et al., 2011). Additionally, during preliminary test sessions, these angles were perceptibly different to both subjects based on measurable differences in their behavioral performances.

Effectively, the VMR perturbed the relationship between neural population activity and the cursor control signal, which caused the cursor to move in an angled manner relative to what would be expected under the baseline decoder (**Figure 1C**). For example, during baseline trials, we expected to consistently observe relatively straight cursor trajectories to the peripheral target. In contrast, we expected to observe significantly curved trajectories during trials immediately following the perturbation (**Figure 1C**). By the end of the perturbation block, we expected to observe cursor trajectories become straighter due to adaptation but still arched relative to baseline (**Figure 1C**).

### Behavioral performance improves rapidly but sub-optimally

We first investigated how the subjects altered their behavior as a result of the VMR perturbation. As expected, during baseline trials, trajectories formed a linear path from the center target to peripheral targets (**Figure 2A – *Baseline***). Conversely, at the onset of the perturbation, trajectories curved (**Figure 2A – *Early Perturbation***). By the end of the perturbation block, trajectories were noticeably less arched (**Figure 2A – *Late Perturbation***).

**Figure 2.**
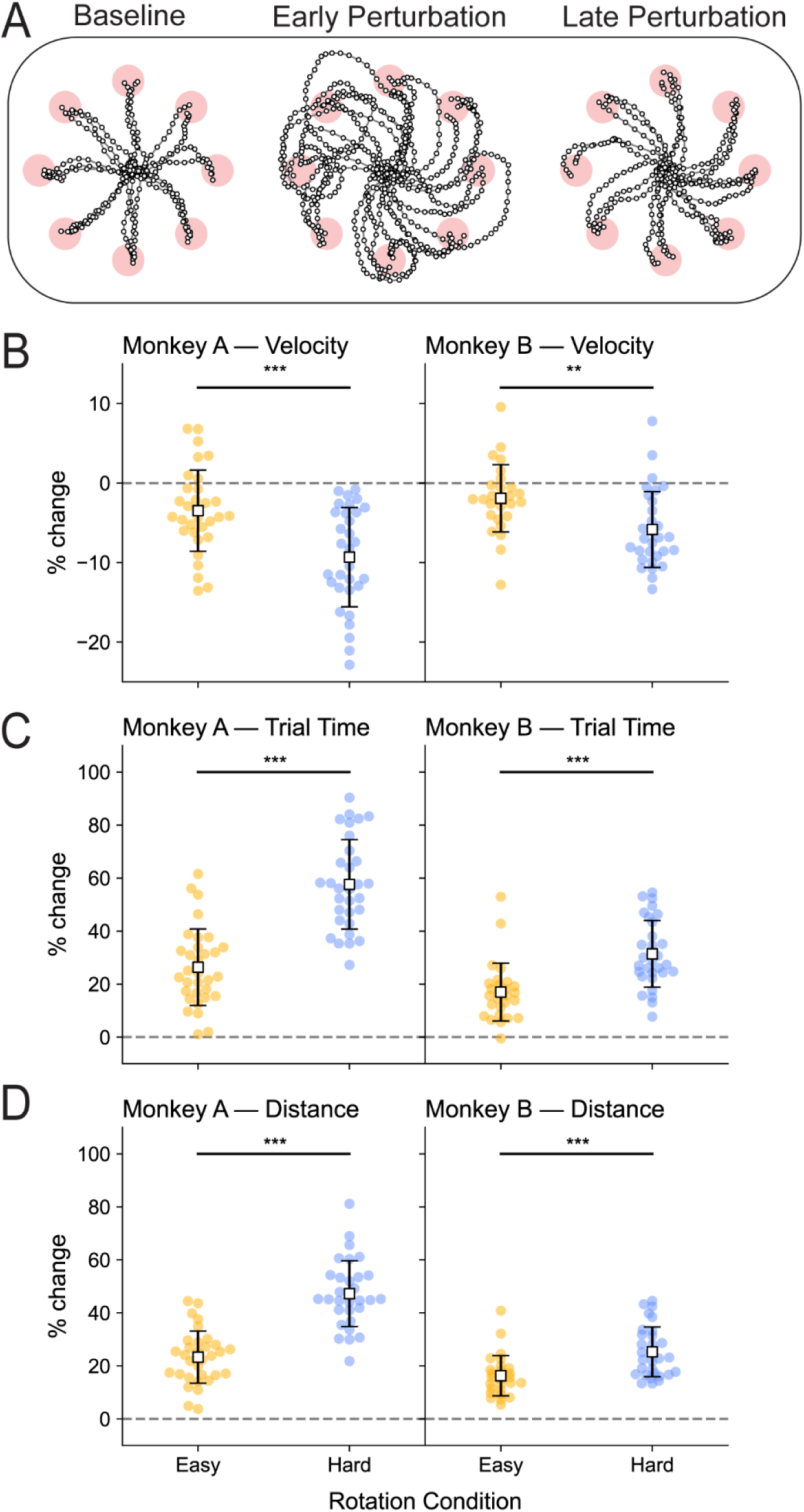
Perturbation leads to worse behavior under the hard condition. (A) Example cursor trajectories to each target (red circles with individual trials connected by grey lines) during baseline, early, and late adaptation, from a representative hard session (Monkey B). Each white scatter point represents a superimposed cursor position update. Only the current cursor position was visible to the subjects. (B) Percentage change in average trial velocity from baseline to perturbation block. Velocity decreased from the baseline to perturbation block for both the easy condition (Monkey A: t_31_ = -3.79, p=6.5e-04; Monkey B: t_26_ = -2.33, p=2.8e-02; two-sided one-sample t-tests comparing to 0% change) and the hard condition (Monkey A: t_30_ = -8.18, p=3.9e-09; Monkey B: t_29_ = -6.61, p=3.0e-07; two-sided one-sample t-tests comparing to 0% change). The decrease in velocity was greater for the hard condition (Monkey A: t_61_ = -4.01, p=1.7e-04; Monkey B: t_55_ = -3.22, p=2.2e-03; two-sided two-sample t-tests). (C) Percentage change in average trial time from baseline to perturbation block. Trial time increased from the baseline to perturbation block for both the easy condition (Monkey A: t_31_ = 10.16, p=2.2e-11; Monkey B: t_26_ = 7.98, p=1.9e-08; two-sided one-sample t-tests comparing to 0% change) and the hard condition (Monkey A: t_30_ = 18.71, p=4.3e-18; Monkey B: t_29_ = 13.50 p=4.9e-14; two-sided one-sample t-tests comparing to 0% change). The increase in trial time was greater for the hard condition (Monkey A: t_61_ = - 7.78, p=1.1e-10; Monkey B: t_55_ = -4.54, p=3.2e-05; two-sided two-sample t-tests). (D) Percentage change in average cursor path length (distance) from baseline to perturbation block. Distance increased from the baseline to perturbation block for both the easy condition (Monkey A: t_31_ = 13.18, p=3.0e-14; Monkey B: t_26_ = 10.95, p=3.1e-11; two-sided one-sample t-tests comparing to 0% change) and the hard condition (Monkey A: t_30_ = 20.84, p=2.1e-19; Monkey B: t_29_ = 14.50 p=8.0e-15; two-sided one-sample t-tests comparing to 0% change). The increase in distance was greater for the hard condition (Monkey A: t_61_ = -8.35, p=1.1e-11; Monkey B: t_55_ = -3.89, p=2.7e-04; two-sided two-sample t-tests). Each scatter point represents a session (Monkey A, easy: n=32, hard: n=31; Monkey B, easy: n=27, hard: n=30). The mean within each rotation condition is represented by a white square with a black outline. The lines extending from the mean are the standard deviation within condition. The easy rotation condition results are presented in yellow, and the hard rotation condition results are presented in blue.

To quantify this observation, we first determined how common behavioral performance metrics (magnitude of velocity, trial time, and distance) changed from the baseline to the perturbation block. Mean velocity decreased from the baseline to perturbation block for both the easy and hard conditions (**Figure 2B**). This corresponded to a worsening in performance: an increase in trial time (**Figure 2C**) and increase in distance (**Figure 2D**). The decrease in performance (i.e., decreased velocity, increased trial time, increased distance) was worse for the hard condition (**Figure 2B-D**).

To quantify this observation on a finer timescale (i.e., sets of trials), we devised a single behavioral performance metric referred to as “performance” (see *Methods – Behavioral Metric*). Performance succinctly combines trial time and distance. These metrics represent the components of velocity, which the subjects directly control through the decoder. In general, trial time and distance are highly related, with short distances generally corresponding to shorter trial times and better behavioral performance. However, to prevent outlier cases in which these two metrics do not agree and to provide a more robust understanding of the behavior of the subjects, we defined a single metric to cohesively represent these metrics. Performance over sets of trials followed an exponential trend (**Figure 3A**). Under both rotation conditions, performance initially increased rapidly and quickly reached an asymptote below baseline performance. These characteristics — rapid improvements and suboptimal adaptation — are consistent with NHP (Jarosiewicz et al., 2008; Chase et al., 2012; Vyas et al., 2018) and human (Krakauer et al., 2000, 2004, 2005, 2019; Tanaka et al., 2009; Huang et al., 2011; Taylor et al., 2014; Kooij et al., 2015; Vaswani et al., 2015; Albert et al., 2021; Matsuda and Abe, 2023) literature.

**Figure 3.**
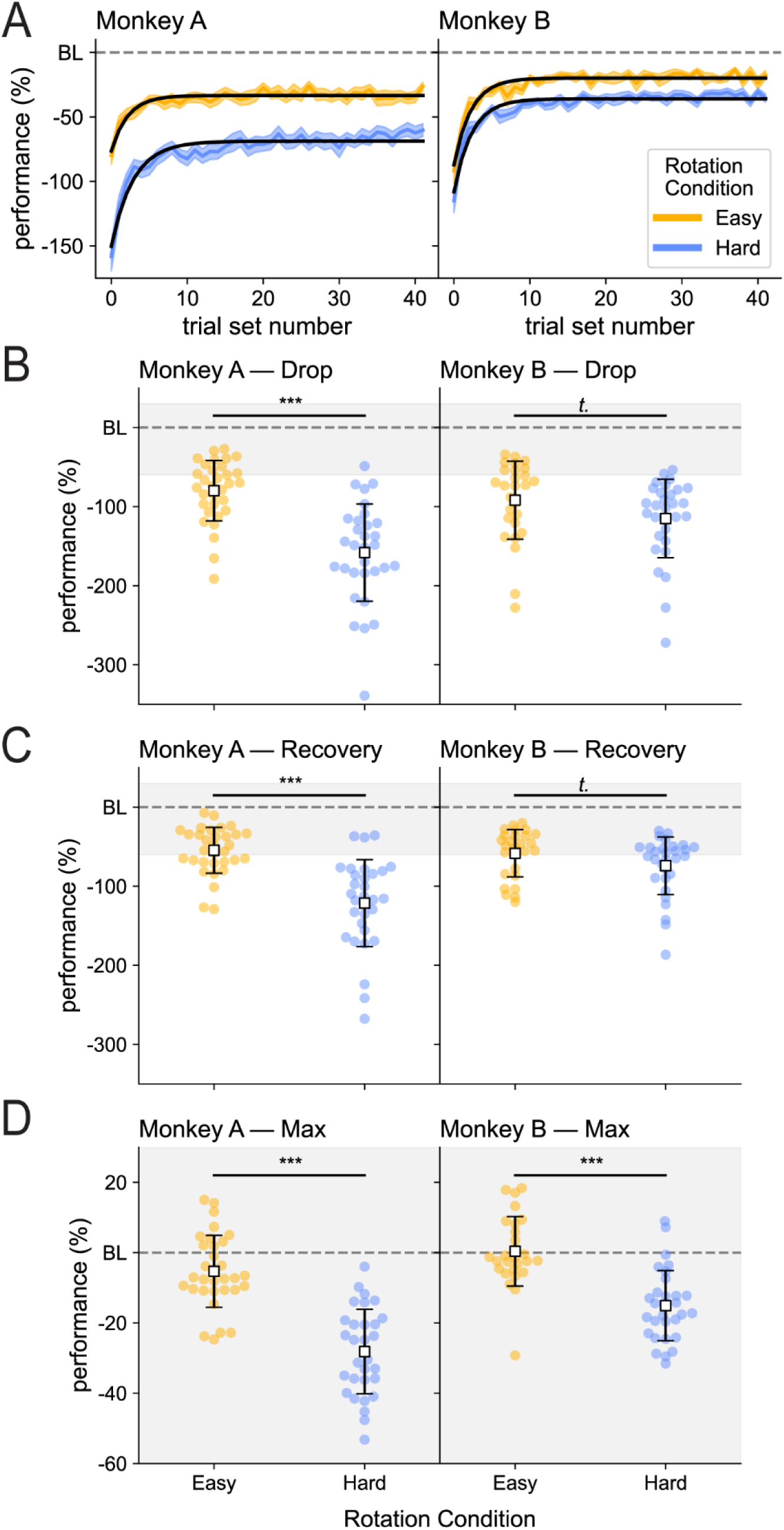
Behavioral adaptation occurs rapidly but sub-optimally. (A) Average performance relative to the baseline block (BL) over sets of eight trials in order of trials sets (TSN: trial set number). Subjects rapidly regained proficient control as indicated by an exponential increase. Qualitatively, subjects achieved higher levels of performance for the easy (yellow) versus hard (blue) rotation condition. However, neither reached an asymptote at baseline (BL) levels. Results presented as mean ± SEM over sessions (Monkey A, easy: n=32, hard: n=31; Monkey B, easy: n=27, hard :n=30). Exponential curves were fit (black line) to the mean (Monkey A, easy: −43.01*e*^−0.46*x*^ − 33.36, R²= 0.9345, hard: − 81.46*e*^−0.35*x*^ − 68.81, R²= 0.9513; Monkey B, easy: −67.26*e*^−0.44*x*^ − 19.82, R²= 0.9593, hard: −72.12*e*^−0.43*x*^ − 35.95, R²= 0.9588; exponential curve with coefficient of determination). (B) Performance relative to BL during the initial drop timepoint (initial set of eight trials in the perturbation block). Performance decreased relative to baseline for the easy (yellow) rotation condition (Monkey A: t_31_ = -11.64, p=7.7e-13; Monkey B: t_26_ = -9.47, p=6.5e-10; two-sided one-sample t-tests comparing to 0% change) and the hard (blue) condition (Monkey A: t_30_ = -14.09, p=9.1e-15; Monkey B: t_29_ = -12.47, p=3.5e-13; two-sided one-sample t-tests comparing to 0% change). The hard (blue) rotation condition imposed a worse initial performance than the easy (yellow) rotation condition (Monkey A: t_61_ = 5.99, p=1.2e-07; Monkey B: t_55_ = 1.74, p=8.7e-02; two-sided two-sample t-tests). (C) Performance relative to BL during the initial recovery timepoint (second set of eight trials in the perturbation block). Performance remained impacted for both the easy (yellow) rotation condition (Monkey A: t_31_ = -10.48, p=1.0e-11; Monkey B: t_26_ = -9.97, p=2.3e-10; two-sided one-sample t-tests comparing to 0% change) and the hard (blue) condition (Monkey A: t_30_ = -12.10, p=4.5e-13; Monkey B: t_29_ = -10.96, p=8.0e-12; two-sided one-sample t-tests comparing to 0% change), more so for the hard condition (Monkey A: t_61_ = 5.91, p=4.2e-07; Monkey B: t_55_ = 1.75, p=8.6e-02; two-sided two-sample t-tests, Monkey A: Welch’s alternative). (D) Performance during the maximum performance timepoint (any set of eight trials in which behavior was close to baseline (0) or exceeded baseline performance (positive performance values)). Performance was better for the easy (yellow) rotation condition in comparison to the hard (blue) rotation condition (Monkey A: t_61_ = 7.99, p=4.5e-11; Monkey B: t_55_ = 3.85, p=3.9e-07; two-sided two-sample t-tests). However, for Monkey A, the performance during the easy condition was still below baseline performance (t_31_ = -2.89, p=7.0e-03; two-sided one-sample t-test comparing to 0% change). For Monkey B, the performance during the easy condition was not significantly different from baseline (t_26_ = 0.1966, p=8.5e-01; two-sided one-sample t-test comparing to 0% change). On average, both subjects had maximum performances below baseline for the hard rotation condition (Monkey A: t_30_ = -12.82, p=1.0e-13; Monkey B: t_29_ = -8.14, p=5.6e-09; two-sided one-sample t-tests comparing to 0% change). For Monkey A, 10 out of 32 easy rotation condition sessions had greater than baseline maximum performance in comparison to 0 out of 31 hard rotation condition sessions. Similarly, for Monkey B, 10 out of 27 easy rotation condition sessions had greater than baseline maximum performance in comparison to 2 out of 30 hard rotation condition sessions. Each scatter point represents a session (Monkey A, easy: n=32, hard: n=31; Monkey B, easy: n=27, hard: n=30). The mean within each rotation condition is represented by a white square with a black outline. The lines extending from the mean are the standard deviation within condition. The easy rotation condition results are presented in yellow, and the hard rotation condition results are presented in blue. The grey area on (B), (C), and (D) represents the range of performance (y-axis) values plotted in (D).

Based on these qualitative observations, we investigated three key timepoints during the adaptation process: initial drop, initial recovery, and maximum performance. The initial drop timepoint was defined as the first set of 8 trials during the perturbation block, and the initial recovery was defined as the second set of 8 trials during the perturbation block. The maximum performance timepoint varied across sessions. As expected, performance under the hard condition was worse than the easy condition at the initial drop, initial recovery, and maximum performance timepoints (**Figure 3B-D**). Still, the performance at the initial drop and initial recovery timepoints was impacted for both rotation conditions, with levels significantly below baseline (**Figure 3B-C**). Performance at the maximum performance timepoint was significantly below baseline under the hard rotation condition for both subjects, and under the easy rotation condition for Monkey A (**Figure 3D**). (Under the easy rotation condition, performance was not significantly different than baseline for Monkey B; **Figure 3D**). In approximately one-third of easy sessions, subjects exceed baseline performance (Monkey A: 31.3%; Monkey B: 37%), in comparison to a zero to small percentage of hard sessions (Monkey A: 0%, Monkey B: 6.7%).

We might expect that the difference in performance at the maximum performance timepoint was due to the limited number of trials. However, the maximum performance timepoint did not significantly differ between the two conditions, which suggests that additional trials would likely not improve adaptation within-session (Monkey A: *t*_61_=-1.24, p=2.2e-01; Monkey B: *t*_55_=0.61, p=5.5e-01; two-sided two-sample t-test).

While the performance was better under the easy condition in comparison to the hard condition across all key timepoints, the rate of adaptation (the amount of performance change per set of trials) did not significantly differ between the two conditions. The rate of adaptation from the initial drop timepoint to the initial recovery timepoint (Monkey A: *t*_61_=-1.00, p=3.2e-01; Monkey B: *t*_55_=-0.90, p=3.7e-01; two-sided two-sample t-test) and from the initial drop timepoint to the maximum recovery timepoint (Monkey A: *t*_61_=-1.24, p=2.2e-01; Monkey B: *t*_55_=-1.14, p=2.6e-01; two-sided two-sample t-test) were not significantly different between the two conditions.

These behavioral results suggest that the initial drop in performance, as a result of the applied VMR perturbation, impacts the possible amount of adaptation. Under the hard condition, subjects started at a lower level of performance which could only be due to the larger rotation angle. Furthermore, subjects ultimately reached a lower level of adaptation at the maximum performance timepoint under the hard condition. However, the rotation condition did not impact the rate of adaptation or the TSN at which the maximum recovery occurred. Overall, these results demonstrate that starting at a lower level of performance at the initial drop timepoint significantly impacts the amount, but not rate of, maximum performance and that the amount of adaptation within a session is limited.

### Neural coordination, represented as shared variance, was negatively impacted – more so under the hard condition

Using factor analysis (FA) (Thurstone, 1934; Williamson et al., 2016; Athalye et al., 2017; Bittner et al., 2017; Zippi et al., 2022), we estimated neural variability in decoder population spiking activity in the baseline and perturbation blocks. In general, FA parses total population variance into private variance and shared variance components. While private variance captures variability that is unique to each decoder unit, shared variance captures how coordinated spike rate variability is across the population.

The total population variance significantly decreased from baseline for both the easy and the hard rotation conditions (**Figure 4A**). This decrease was significantly greater under the hard condition (**Figure 4A**). Similarly, the amount of shared variance decreased from the from baseline under both rotation conditions, more so for under the hard condition (**Figure 4B**). While private variance did decrease significantly under the hard condition for both subjects, it only decreased significantly under the easy condition for Monkey B (**Figure 4C**). Furthermore, there was not a significant difference between conditions (**Figure 4C**). Therefore, the difference in decrease in total variance, that separates the two conditions, is due to changes in shared variance. Overall, the perturbation disrupted neural population coordination – more so for the more difficult condition.

**Figure 4.**
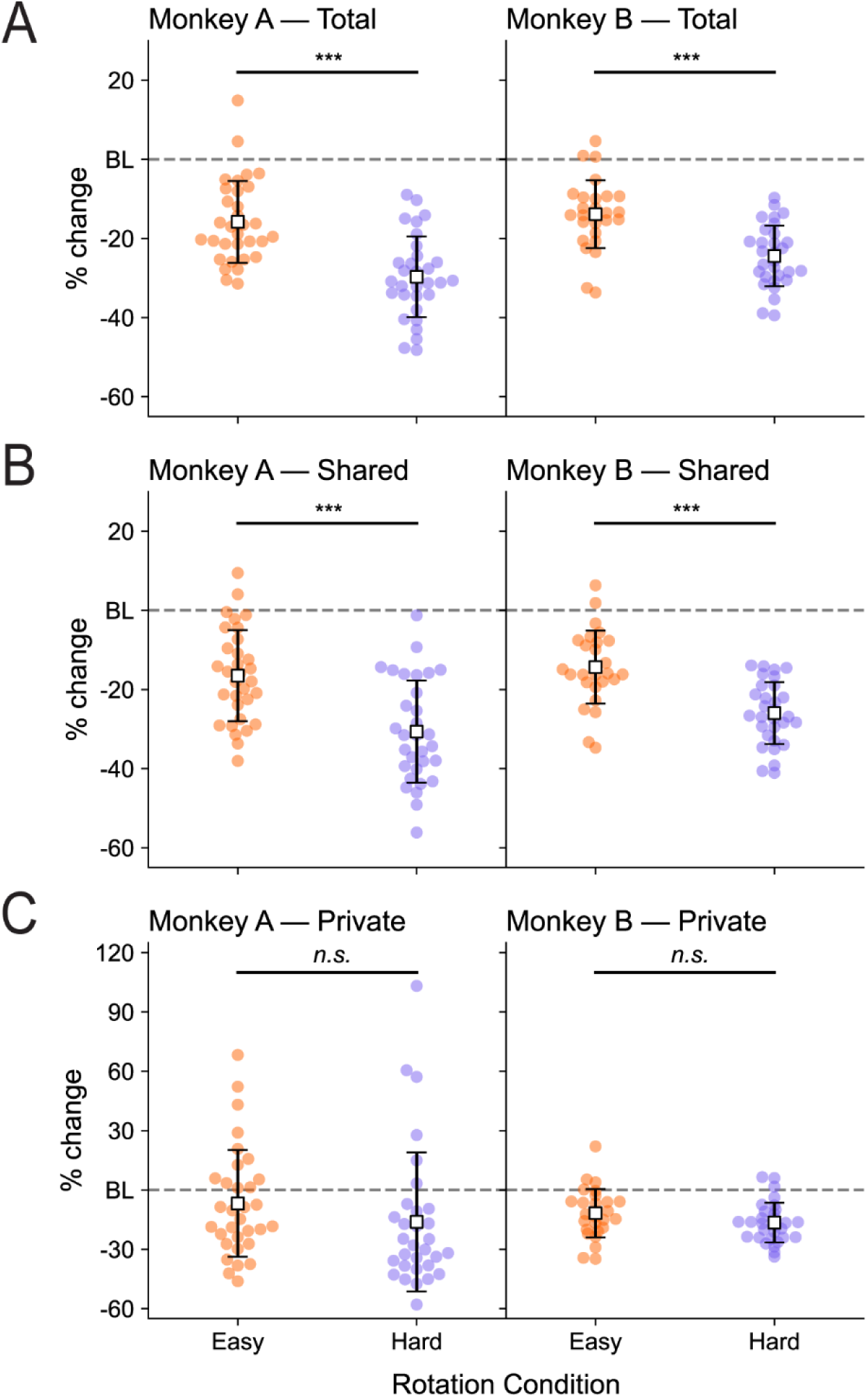
Neural coordination, represented as shared variance, was negatively impacted – more so for under the hard condition. (A) Percent change from the baseline (BL) block to the perturbation block in total amount of variance explained by FA models. Total variance is the sum of shared and private variance. Total variance decreased for both the easy (orange) rotation condition (Monkey A: t_31_ = -8.52, p=1.3e-09; Monkey B: t_26_ = -8.24, p=1.30e-08; two-sided one-sample t-tests comparing to 0% change) and the hard (purple) condition (Monkey A: t_30_ = -15.98, p=3.2e-16; Monkey B: t_29_ = - 17.23, p=9.0e-17; two-sided one-sample t-tests comparing to 0% change) – more so for the hard rotation condition (Monkey A: t_61_ = 5.29, p=1.8e-06; Monkey B: t_55_ = 4.83, p=1.1e-05; two-sided two-sample t-tests). (B) Percent change from the baseline (BL) block to the perturbation block in the amount of shared variance explained by FA models. Shared variance decreased for both the easy (orange) rotation condition (Monkey A: t_31_ = -7.97, p=5.3e-09; Monkey B: t_26_ = -7.93, p=2.1e-08; two-sided one-sample t-tests comparing to 0% change) and the hard (purple) condition (Monkey A: t_30_ = -13.00, p=7.3e-14; Monkey B: t_29_ = -17.84, p=3.6e-17; two-sided one-sample t-tests comparing to 0% change).– more so for the hard rotation condition (Monkey A: t_61_ = 4.53, p=2.8e-05; Monkey B: t_55_ = 5.05, p=5.2e-06; two-sided two-sample t-tests). (C) Percent change from the baseline (BL) block to the perturbation block in the amount of private variance explained by FA models. Private variance decreased significantly under the easy (orange) rotation condition for Monkey B only (Monkey A: *t*_31_=-1.40, p=1.7e-01; Monkey B: *t*_26_;= -4.87, p=4.7e-05 two-sided one-sample t-tests comparing to 0% change). Private variance decreased significantly under the hard (purple) rotation condition for both subjects (Monkey A: t_30_ = -2.51, p=1.8e-02; Monkey B: t_29_ = -8.88, p=9.1e-10; two-sided one-sample t-tests comparing to 0% change). There was not a significant difference between the conditions (Monkey A: t_61_ = 1.17, p=2.5e-01; Monkey B: t_55_ = 1.60, p=1.2e-01; two-sided two-sample t-tests). Each scatter point represents a session (Monkey A, easy: n=32, hard: n=31; Monkey B, easy: n=27, hard: n=30). The mean within each rotation condition is represented by a white square with a black outline. The lines extending from the mean are the standard deviation within condition. The easy rotation condition results are presented in orange, and the hard rotation condition results are presented in purple.

One additional consideration in interpreting these results is changes in mean spike rates. Larger mean spike rates generally correspond to higher amounts of variability, following a Poisson or super-Poisson mean-to-variance relationship. Importantly, these results were not driven by changes in firing rates, as the average firing rates of decoder units did not change statistically significantly from baseline to perturbation for any session for either subject.

### Dimensionality does not change significantly but the similarity of factors across before and after perturbation does

The number of factors identified by FA did not change significantly from the baseline block model to the perturbation block model under the easy (Monkey A: t_31_ = 1.19, p=2.4e-01; Monkey B: t_26_ = 0.76, p=4.5e-01; two-sided one-sample t-tests comparing to 0% change) or the hard condition (Monkey A: t_30_ = 1.01, p=3.2e-01; Monkey B: t_29_ = -1.38, p=1.77e-01; two-sided one-sample t-tests comparing to 0% change), and did not differ between conditions (Monkey A: t_61_ = -0.05, p=9.6e-01; Monkey B: t_55_ = 1.50, p=1.4e-01; two-sided two-sample t-tests).This is aligned with our expectations that the estimated dimensionality for the intrinsic manifold would not change significantly on the short timescale of adaptation. Units are more likely to jointly reassociate activity to counteract the perturbation than to develop novel dimensions.

We then identified “major” and “minor” factors by sorting the factors by the amount of shared variance explained. The first major factor explained approximately 50% of the shared variance (Monkey A: *m*_*baseline*_ =48.46%±0.77%, *m*_*perturbation*_ =47.30%±0.69%; Monkey B: *m*_*baseline*_ = 53.55%±0.83%, *m*_*perturbation*_ =53.15%±0.78%; mean ± standard error of the mean), and the second major factor explained approximately 30% of the shared variance (Monkey A: *m*_*baseline*_=31.28%±0.71%, *m*_*perturbation*_=31.77%±0.80%; Monkey B: *m*_*baseline*_= 29.65%±0.79%, *m*_*perturbation*_=31.47%±0.69%; mean ± standard error of the mean). Interestingly, we found that the two major factors were related more strongly across blocks than the remaining minor factors were amongst themselves (**Figure 5**). Additionally, the strength of the relationship was greater for the easy rotation condition.

**Figure 5.**
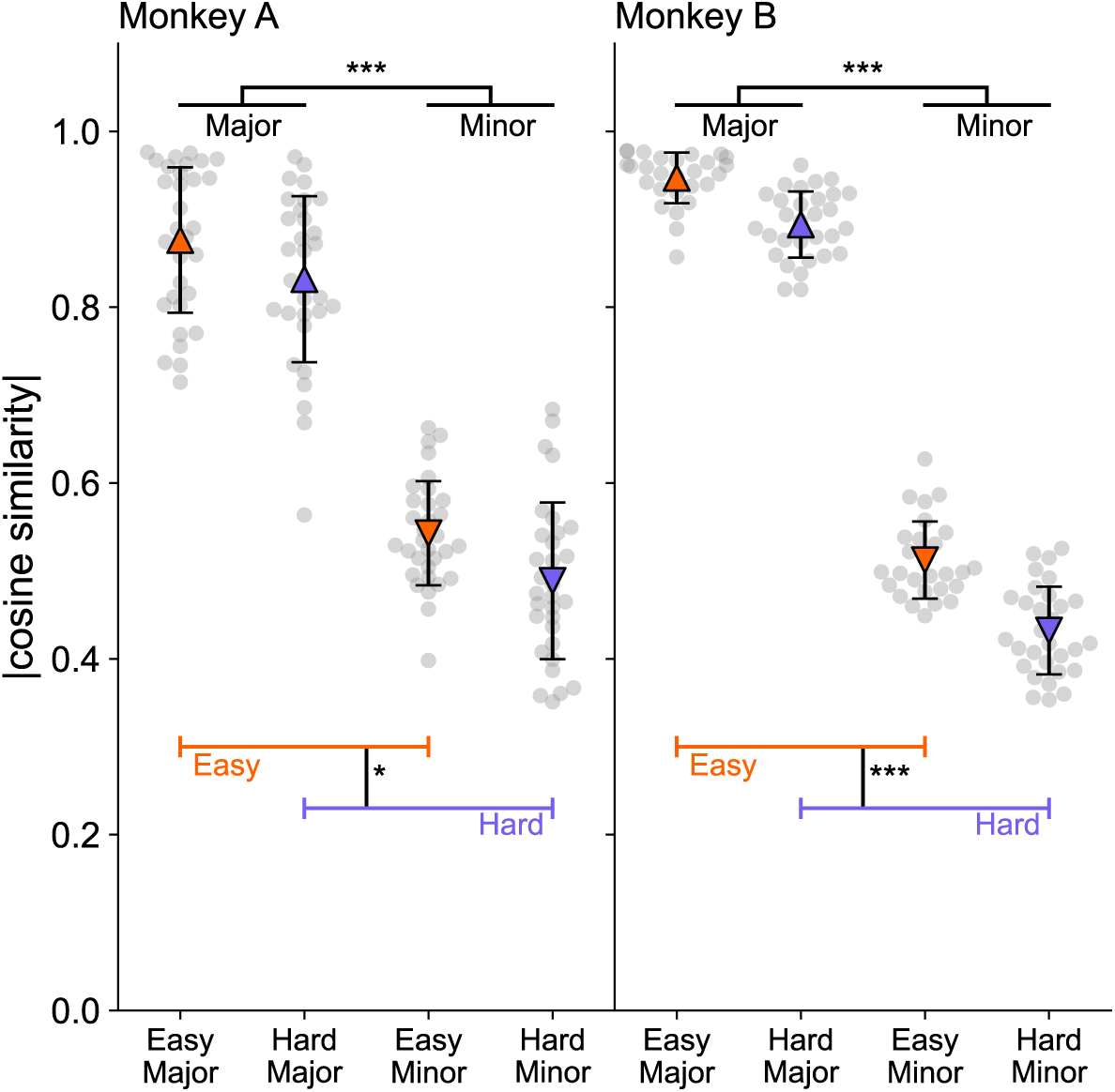
Major factors are largely maintained after perturbation while minor factors are less so. The relationship of factors across blocks was measured as the absolute value of cosine similarity between pairs of factors (one baseline factor, one perturbation factor). Perturbation major factors were only compared to baseline major factors, and perturbation minor factors were only compare to baseline minor factors. For major factors, we determined, of the major factors, which value maximized the absolute value of cosine similarly and averaged the results together. We computed the average absolute cosine similarity value for minor factors in a similar fashion, only comparing minor factors from perturbation to minor factors in baseline. A two-way ANOVA with factors of rotation condition (levels: 50°, 90°) and classification (levels: major, minor) revealed significant main effects for both rotation condition (Monkey A: *F*_1,122_= 10.98, p=1.2e-03; Monkey B: *F*_1,110_ =72.37, p=9.9e-14) and classification (Monkey A: *F*_1,122_ =515.74, p=1.2e-45; Monkey B: *F*_1,110_ =3302.56, p=7.0e-84). There was no significant interaction effect for Monkey A (*F*_1,122_ =0.10, p=7.5e-01) and a trending interaction effect for Monkey B (*F*_1,110_ =0.005, p=8.82e-02).

Taken together, these results suggest that neural populations largely maintain underlying population activity structure and alter modes of co-modulation that constitute less shared variance, highlighting a neural limitation in adaptation. Because the factors were overall more similar under the easy condition than under the hard condition, the difficulty condition may impose the need for larger changes to modes of co-modulation which are harder to develop on a short timescale. Through the adaptation process, subjects were able to alter neural activity and, thus, their behavior to largely counteract the perturbation. However, the cursor trajectories were still curved relative to baseline at the end of the session (**Figure 3A**). The subjects likely needed to generate novel patterns (i.e., factors) to optimize performance which takes periods of days (Oby et al., 2019).

### Neural variance highlights variability in behavioral performances

In addition to highlighting differences between difficulty conditions, neural variance metrics correlate to varying behavioral performances we observed across sessions. Changes in shared variance were correlated with the performance at the initial recovery and maximum performance timepoints (**Figure 6A-B**). Conversely, changes in private variance were not reliably (i.e., consistently significant between subjects) correlated with performance at either of these timepoints. Neither metric was reliably correlated with performance at the initial drop timepoint.

**Figure 6.**
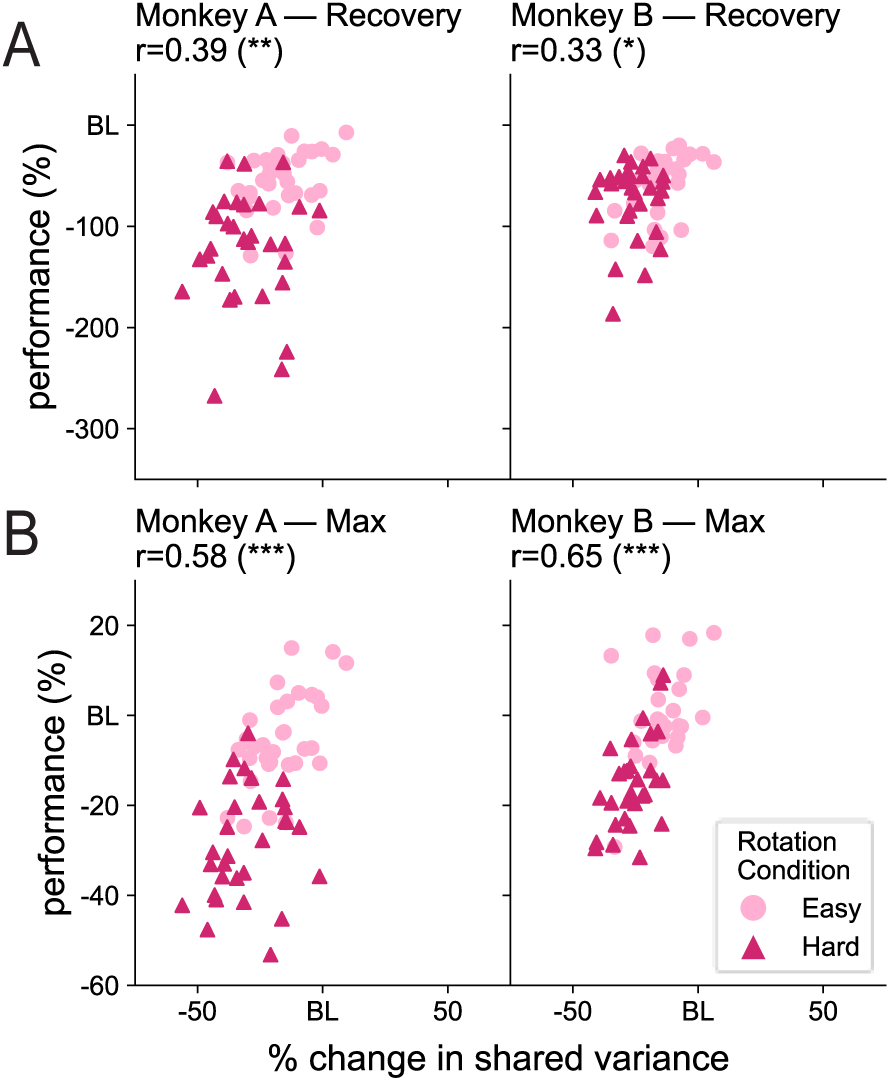
Change in shared variance –but not private variance – correlates with recovery in performance. (A) Larger amounts of shared variance in the perturbation block correlated with better performance at the initial recovery timepoint (Monkey A: r=0.39, p=1.5e-03; Monkey B: r=0.33, p=1.2e-02; Pearson’s correlation coefficient). Not pictured: Changes in private variance were not correlated with performance at the initial recovery timepoint for either subject (Monkey A: r=0.19, p=1.3e-01; Monkey B: r=0.18, p=1.8e-01; Pearson’s correlation coefficient). (B) Larger amounts of shared variance in the perturbation block correlated to better performance at the maximum performance timepoint (Monkey A: r=0.58, p=7.7e-07; Monkey B: r=0.65, p=4.0e-08; Pearson’s correlation coefficient). Changes in private variance were significantly correlated with performance at the maximum performance timepoint for Monkey B (r=0.66, p=32.0e-08; Pearson’s correlation coefficient) but not for Monkey A (r=0.14, p=2.8e-01) Each scatter point represents a session (Monkey A, easy, circle: n=32, hard, triangle: n=31; Monkey B, easy, circle: n=27, hard, triangle: n=30).

The amount of shared variance in the perturbation block was correlated to performance at the initial drop and initial recovery timepoints (**Figure 7A-B**), whereas only private variance in the perturbation block was reliability correlated with the performance at the maximum performance timepoint (**Figure 7C**). Interestingly, higher levels of private variance correlated with better performance, which is counterintuitive to the general belief that private variance corresponds with noise. While the number of units is moderately correlated to the amount of private variance (Monkey A: r=0.43, p=4.3e-04; Monkey B: r=0.63, p=1.8e-07; Pearson’s correlation coefficient), the number of units was not correlated with the performance at the maximum performance timepoint (Monkey A: r=-0.17, p=1.8e-01; Monkey B: r=0.22, p=1.1e-01).

**Figure 7.**
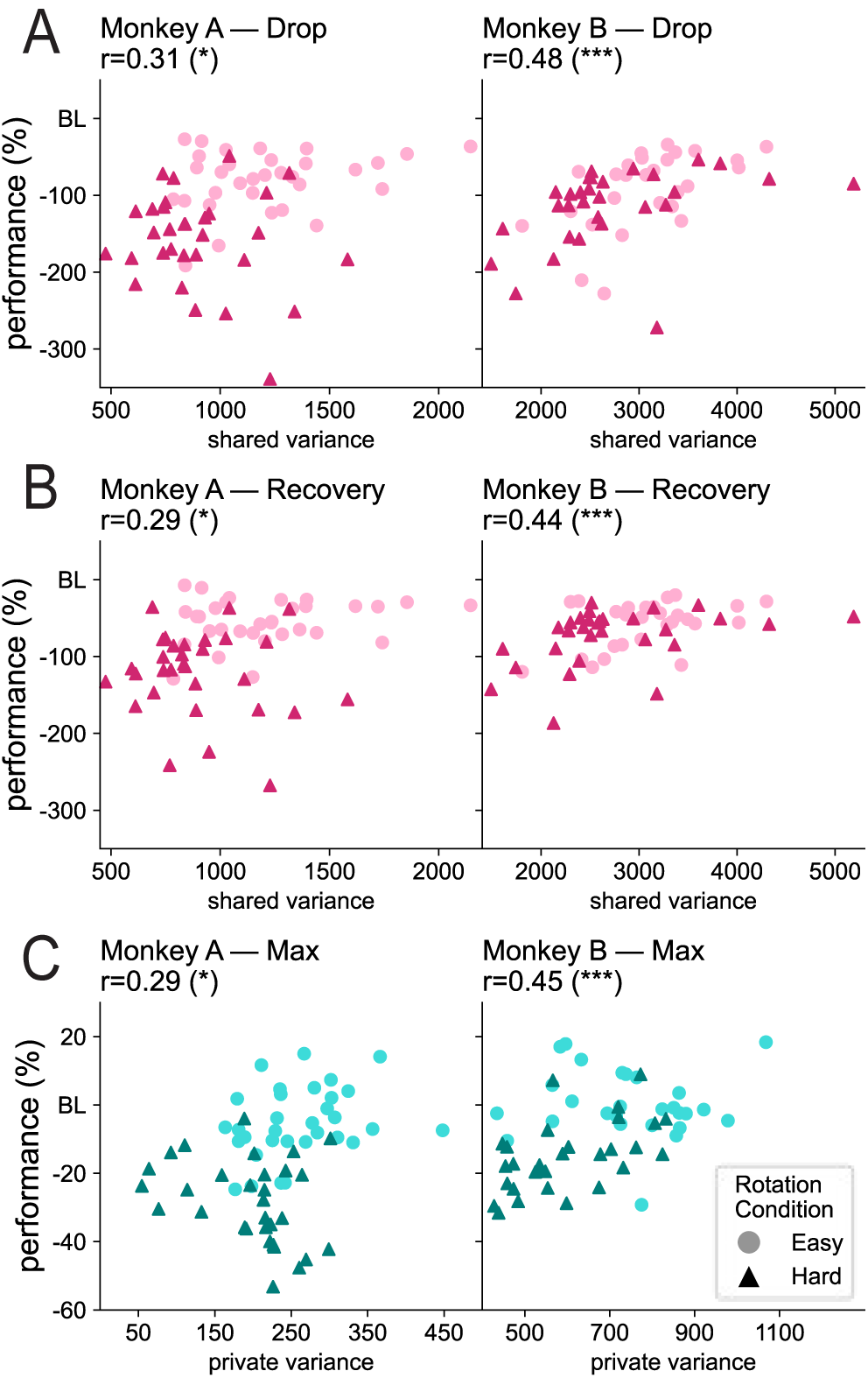
Amount of neural variance predicts behavioral performance at the three key timepoints. (A) Larger amounts of shared variance in the perturbation block at the initial drop timepoint correlated with better performance (Monkey A: r=0.31, p=1.5e-02; Monkey B: r=0.48, p=1.4e-04; Pearson’s correlation coefficient). Not pictured: For Monkey A, private variance in the perturbation block exhibited a trending correlation with performance (r=0.22, p=8.7e-02; Pearson’s correlation coefficient), whereas, for Monkey B, the correlation was not significant (r=0.16, p=2.5e-01; Pearson’s correlation coefficient). (B) Larger amounts of shared variance in the perturbation block at the initial recovery timepoint correlated to better performance (Monkey A: r=0.29, p=2.0e-02; Monkey B: r=0.44, p=6.6e-04; Pearson’s correlation coefficient). Not pictured: For Monkey A, private variance in the perturbation block had a significant correlation with performance (r=0.27, p=3.3e-02; Pearson’s correlation coefficient), but the correlation was not significant for Monkey B (r=0.21, p=1.1e-01; Pearson’s correlation coefficient). (C) Larger amounts of private variance in the perturbation block at the maximum performance timepoint correlated to better maximum performance (Monkey A: r=0.29, p=1.9e-02; Monkey B: r=0.45, p=4.2e-04; Pearson’s correlation coefficient). Not pictured: For Monkey A, shared variance in the perturbation block was significantly correlated with performance (r=0.41, p=8.7e-04; Pearson’s correlation coefficient), but the correlation was not significant for Monkey B (r=0.03, p=8.0e-01; Pearson’s correlation coefficient). Each scatter point represents a session (Monkey A, easy, circle: n=32, hard, triangle: n=31; Monkey B, easy, circle: n=27, hard, triangle: n=30).

Baseline variance metrics were not reliably predictive of the performance. This suggests that the relationship between amounts of variance is specific to the adaptation process.

## Discussion

Here, we demonstrated that measures of neural population variance are positively and significantly correlated with the behavioral adaptation process in addition to highlighting differences and limitations in adaptation between difficulty conditions and across sessions Identifying. We investigated this relationship through using a linear KF to mathematically define the mapping between neural spiking activity and computer cursor movement. We then perturbed the mapping by jointly rotating the neural decoder unit activity to study how decoder unit activity patterns changed. Importantly, we did not impose any constraints on the strategy subjects used to modulate variability in their neural activity. Therefore, any correlations between neural variance and behavior are outside of the constraints of the causal relationship established through the BCI.

### The role of neural variance in balancing stable and flexible behavior

Both easy and hard rotation conditions imposed the same type of perturbation to neural patterns: neural activity is altered jointly. Essentially, this requires that subjects adapt existing neural patterns to “new axes” in the neural space. Given the evidence presented here, the rate of adaptation is not impacted by perturbation difficulty; however, the amount of coordinated neural activity is. Therefore, adapting to the easy and hard rotations likely involves similar neural strategies in which harder perturbations limit the maximum amount of adaptation through causing a greater disruption in the initial amount of shared variability.

Previously, other groups have investigated VMR-BCI tasks in NHPs (Jarosiewicz et al., 2008; Chase et al., 2012; Vyas et al., 2018). Notably, neural units employ a seemingly sub-optimal mixture of single-decoder-unit activity and population-wide strategies to re-associate existing patterns to meet task demands. Employing a sub-optimal strategy appears to highlight a limitation. However, while the optimal solution for a given task may be to completely re-orient activity to the new frame of reference, factors such as perceived error magnitude (Marko et al., 2012; Matsuda and Abe, 2023), importance of error signal to the task at hand (Wei and Körding, 2009), and noise in feedback (Vaswani et al., 2015; Albert et al., 2021) likely place a constraint on how much a subject should adapt (e.g., not updating when environmental cues are noisy) to keep a balance between stable behavior and flexible adaptation. Similarly, from our work presented here, we observed private variance in both the learned baseline behavior and throughout the adaptation process. Based on our definition of total variance, this suggests that decoder populations could further achieve adaptation back to baseline levels or improve baseline performance via increasing shared variance (i.e., decreasing private variance). Recent evidence, however, has suggested that private variance is likely not pure noise but, instead, enables exploration (Athalye et al., 2017; Sternad, 2018) and adaptation. As demonstrated in **Figure 7C**, private variance had a significant, positive correlative relationship to behavioral performance. Still, as research in humans has suggested, excessive variability in neural activity could be indicative of maladaptive brain states (Murtha et al., 2002; Williams et al., 2005; MacDonald et al., 2006; Nomi et al., 2018; Waschke et al., 2021).Therefore, investigating the role of neural variability in this balancing act, as performed here, is necessary to gain insight into both healthy and pathological activity.

### Limitations of the study

While this study has contributed to the growing body of work that highlights the limitations of neurofeedback adaptation and learning, we would like to highlight some limitations that would have further strengthened our results. We did our due diligence to demonstrate that the behavioral results are not idiosyncratic to one animal, and two subjects is a common number in primate research; however, we conceded that our behavioral results could only be strengthened by including additional subjects. Furthermore, by adding additional subjects, we would be able to include females and investigate any possible effects of biological sex as a variable. Additionally, we sampled a limited number of rotation conditions. While this was sufficient for demonstrating the neural differences in behavior for “within-manifold” perturbations, we believe that our later analyses (**Figure 6**, **Figure 7**) could benefit from additional sampling of neural population variance changes to provide a more comprehensive view of the predictive relationship of neural population variance and behavior. Finally, although subjects routinely performed hundreds more trials than what was analyzed here, we did not directly measure engagement with the task. Had we measured external factors that could explain changes in variability (e.g., attention), we could further parse neural population changes driven by changes in task demands versus neural population changes driven by task-irrelevant sources. Regardless of these limitations, our work still presents significant relationships between neural and behavioral features of adaptation.

### Future directions in modeling

The exponential profiles of behavioral adaptation (**Figure 3B**) are qualitatively similar in profile to the human behavioral models: rapid improvements (Krakauer et al., 2000, 2004, 2005; Tanaka et al., 2009; Huang et al., 2011; Taylor et al., 2014) are followed by sub-optimal adaptation. (Kooij et al., 2015; Vaswani et al., 2015; Albert et al., 2021; Matsuda and Abe, 2023) While human behavioral models have successfully captured the tradeoff between stability and flexibility through examining contributions of implicit and explicit strategies (Krakauer et al., 2000, 2019; Huang et al., 2011; Taylor et al., 2014; Albert et al., 2021), these models do not seek to explain the underlying neural mechanisms. Here, we presented critical neural correlates that explain differences in key behavioral adaptation timepoints.

Our behavioral and neural profile can be divided into multiple components based on producing beneficial output for a task goal. When estimating components of neural variability, the FA model has no knowledge of task states. Therefore, the variance captured is not necessarily all task-related (e.g., attentional shifts); however, all patterns were read out in the task space. While not all neural variability is productive to the task goal, subjects do receive feedback on any undesired errors this causes. To fully compare our results with the human behavioral models, further investigation into specific explicit and implicit strategies as well as influences of non-task-related variables on neural variability is needed.

### Implications for neurofeedback paradigms

The ability to explore limitations in self-regulated neurofeedback with neural ensemble resolution is an important complementary body of research that is needed to improve neurofeedback paradigms (Sitaram et al., 2017; Lubianiker et al., 2022).The use of BCIs, and more broadly neurofeedback training, has recently increased in the clinical realm in applications such as a tool to better understand autism (Dinstein et al., 2015), stroke rehabilitation (Mane et al., 2020; Ganguly et al., 2022) and attention deficit hyperactivity disorder (ADHD) focus training (Chabot et al., 2001; Enriquez-Geppert et al., 2019). Human studies involving simultaneous behavior and imaging have provided critical evidence in identifying neural substrates and networks for clinical neurofeedback applications in which patients endogenously alter their brain activity to a desired state. However, the underlying mechanisms at the neural level are not well characterized (Sitaram et al., 2017; Lubianiker et al., 2022). This, in part, has led to clinical variability in the effectiveness of this paradigm (Sitaram et al., 2017; Lubianiker et al., 2022). Ultimately, through identifying a neural framework in which adaptation can be tracked and predicted, we have begun to identify critical limitations of neurofeedback training.

## Materials and Methods

### Code accessibility

All experiments and analyses were run using Python3 (version 3.9.7). Original experimental code (BMI3D library) can be found here: https://github.com/carmenalab/brain-python-interface, and customized experimental code can be found here: https://github.com/santacruzlab/bmi_python. Analysis scripts are available upon request.

### Experimental model: Rhesus macaque

Two group-housed, healthy male rhesus macaques (*Macaca mulatta*) were used to perform this study (Monkey A: age 5 years, ∼9.5 kg; Monkey B: age 5 years, ∼8.8 kg). All rhesus macaque procedures were approved by the Institutional Animal Care and Use Committee (IACUC) of The University of Texas at Austin, a fully AAALAC-accredited institution. The experiments were conducted in accordance with the *Guide for the Care and Use of Laboratory Animals*, Public Health Service Policy, and the Animal Welfare Act and Regulations. Prior to any procedures, both subjects underwent MRI brain scans (Siemens Skyra 3.0T MRI housed at the University of Texas at Austin) to assist with neural targeting. To ensure stability of the recordings and engagement in the task, subjects were then implanted with a headpost to constrain their head during behavior. We permitted six weeks for adequate bone integration prior to use of the headpost and chronic microelectrode array (MEA) implantation. We implanted tungsten (35 µM) MEAs (Innovative Neurophysiology, Inc. Durham NC) into the left hemisphere primary motor (M1) / pre-motor (PMd) area of cortex for both subjects (Monkey A: 16x4 electrodes, Monkey B: 16x8 electrodes). Using an atlas of stereotaxic coordinates(Saleem and Logothetis, 2006) and MRI-based brain models, we estimated the coordinates for the motor cortex. In surgery, the location of the MEA insertion was visually confirmed based on topological cerebral landmarks.

### Electrophysiology

We simultaneously recorded behavioral task messages and continuous (local field potential) and discrete (spike) signals using the Grapevine NIP (Ripple Neuro, Salt Lake City, UT). Local field potential signals were not used in this analysis. Prior to each session, putative spiking units were sorted using a hooping method (Ripple Neuro, Salt Lake City, UT). Briefly, we defined each waveform using one to four threshold ranges. A “spike” was considered the timestamp of the start of a waveform event that crossed into all defined hoops. Distinctions between single- and multi-unit activity were not made and subsequently all treated as units forming the decoder input.

### Behavioral training

Prior to instantiating the BCI task, subjects were trained on the same center-out task using a joystick (manual control). Subjects sat in a specialized primate chair (Crist instrument, Hagerstown, MD) approximately 18” (∼46 cm) from a 24” computer monitor. The manual control task took approximately 1-2 months to learn. Post joystick training, transfer to the BCI task occurred within the first session. Critically, this was due to having the subjects perform a KF decoder “tuning” session (see *Kalman Filter Decoder*) prior to the start of the main VMR task each session.

Subjects were allowed unlimited attempts to complete a trial (successful hold of a peripheral target), and only successful attempts were analyzed in this script. Therefore, each trial set contains one trial to each of the eight targets (in a pseudorandom order). In general, the subjects were able to complete the vast majority of trials within one to two attempts.

To maintain task engagement, subjects were acclimated to a regulated fluid intake protocol and head restraint during the task in accordance with our approved IACUC protocol. The amount of fluid earned per day was dependent on the number of trials completed. Subjects typically worked until they were satiated, and the session was stopped when the subject did not move the cursor for more than five minutes. A constant cursor size of radius 0.5 cm and target radius size of 2 cm were used.

To run the manual control and BCI tasks, we adapted publically available code from a Pythonic library named “BMI3D” that was originally developed through the Carmena Lab at the University of California, Berkeley. In addition to updating cursor movements on a computer screen, the task code transmitted messages regarding task state at a rate of 60 Hz to an Arduino board. Messages and timestamps were recorded as analog inputs to the acquisition system and used offline to synchronize behavioral and neural data. This board was also used to trigger a liquid reward by sending a TTL signal to a solenoid-based system that would release approximately 100µL of juice per reward period.

### Behavioral metrics

In **Figure 2**, we examined how common behavioral metrics (magnitude of velocity, trial time, distance) changed from the baseline block to the perturbation block. The magnitude of velocity was computed prior to averaging across all trials in a block. Trial time was the average time elapsed from the onset of cursor movement to the start of the peripheral hold period. Distance was computed as the average pathlength that the cursor travelled during a trial. Percent change for each metric during each session was computed as the difference between perturbation behavior and baseline behavior, divided by the baseline behavior, and multiplied by 100%.

To examine behavior on a finer timescale (i.e. non-overlapping sets of 8 trials) in **Figure 3**, we consolidated the trial time and distance metrics in a single metric termed “Performance.” The average trial times and distances were computed for each set of 8 trials in the perturbation block. Then, the percent change from the respective mean baseline metrics were calculated. This normalization ensured that trial time and distance were on the same scale. For each set of trials, we represented the trial time and distance as a single point in two-dimensional Cartesian space. Additionally, by these conventions, baseline performance was (0,0). “Performance” was calculated as the straight-line distance between baseline and each pair of metrics for each set of trials. To aid in intuition, a correction factor or -1 was multiplied to the distances if either or both of the individual metrics were greater than baseline (i.e., worse performance). Negative performance values indicate performance lower than BL, while positive values indicate performance exceeding BL.

### Kalman filter decoder

The Kalman Filter (KF) is commonly used to linearly map spike vector data to computer cursor velocity(Wu et al., 2006; Dangi et al., 2014; Sadtler et al., 2014). The model consists of two key equations: the state-space model of the cursor dynamics (**Equation 1**) and the neural observation model (**Equation 5**).

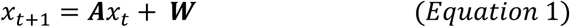

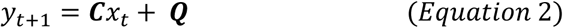

The models are parameterized by the set of matrices {**A**, **C**, **W**, **Q**}. **A** is a matrix of constants that describes how intended velocity evolves with time. **C** relates intended velocity to the spike count vector (**Y**). **W** and **Q** are additive Gaussian noise terms. **A** remains constant over all experimental sessions and is not impacted by neural input. Once the KF has been trained, the resultant model can be expressed as a steady-state, linear, time-invariant form as in **Equation 3**, with **x** being a vector containing the horizonal and vertical components of cursor movement (**Equation 4**):

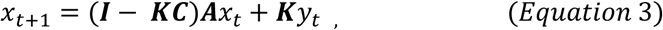

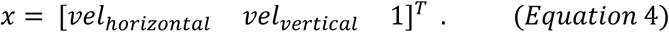

For each decoder unit, a separate Kalman gain (**K**) is applied for vertical and horizontal velocity components. Position is determined from velocity by adding the previous position plus velocity times the length of a decoder update (100 ms; **Equation 5**):

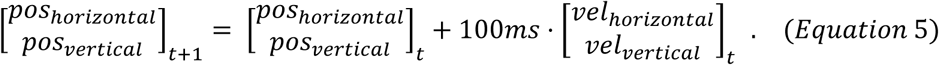

To induce a VMR perturbation, we applied a counterclockwise rotation matrix (**Equation 6**) that altered the gain (**K**) for each decoder unit (**Equation 7**):

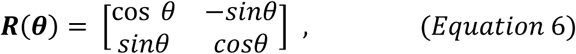

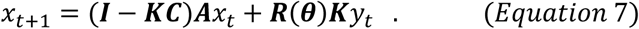

At the start of each experimental session (day), subjects passively observed a cursor moving from the center of the screen to one of eight possible peripheral targets and then back to center screen (visual feedback task, VFB). Data from this task (spiking data from 6 trials x 8 targets; ∼3.5 minutes) was used to train the initial KF decoder. We selected units that fired at a rate of >1Hz and had an amplitude of >80µV, on average, over the VFB training data. To tune the decoder, we implemented closed-loop decoder adaptation (CLDA) for 5 to 10 minutes using a recursive maximum likelihood function (Dangi et al., 2014). Essentially, CLDA acted as a two-learner system in which cursor movement is a combination of neural input from the subject and straight-line assistance from the computer. Over this timespan, the computer assist would decrease linearly from 10% to 0% assist. CLDA was not applied during the main task.

### Factor analysis

We used FA to decompose neural population activity into mean, shared (correlated), and private (uncorrelated) components. We modeled population activity as in **Equation 8** (Williamson et al., 2016; Athalye et al., 2017; Bittner et al., 2017):

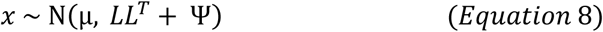

where *x* ∈ ℝ^*N*^ is the spike counts of *N* neurons, *μ* ∈ ℝ^*N*^is a vector of mean spike counts for each neuron (*N*) in the population, *L* ∈ ℝ^*N*×*M*^ represents the factor loading matrix of *M* factors, and Ψ ∈ ℝ^*N*×*N*^ is the private variance matrix. To estimate the model parameters, we used the expectation-maximization (EM) algorithm (Dempster et al., 1977; Williamson et al., 2016; Bittner et al., 2017).

We defined the shared variance, private variance, and total variance explained as in **Equation 9**, **Equation 10**, and **Equation 11**, respectively:

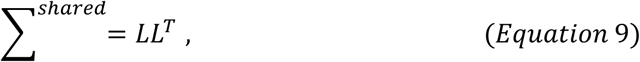

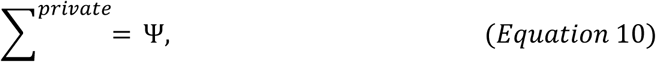

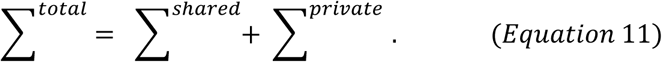

We fit a separate FA model for each baseline and perturbation block. The dimensionality of the input to each model was the number of trials by the number of units. While the number of trials was kept constant (8 targets*42 trials = 336 trials), the number of units varied by session (but not within session). Each entry was the unit’s trial-averaged spike rate (Hz). To determine the number of factors used to fit each model, we performed repeated 10-fold cross-validation (CV) and chose the number of factors that maximized the average log-likelihood score. In practice, we used the *cross_val_score* function from the *scikit-learn* library (version 1.0.1) to perform CV, and the *FactorAnalysis* function from the *scikit-learn* library (version 1.0.1) to fit the FA models.

### Statistical analyses

We performed all statistical and mathematical modeling in custom-written Python 3 scripts using the *SciPy* library (version 1.7.0). The statistical details for each figure can be found in the figure legends. In general, we assumed a significance level of *α* = 0.05. Trending (*t.*) p-values were defined as being greater than 0.05 but less than 0.1, and non-significant (*n.s.*) results were greater than or equal to 0.1. One star (*) indicated a p-value less than or equal to 0.05 and greater than 0.01. Two stars (**) indicated a p-value less than or equal to 0.01 but greater than 0.001. Three stars (***) indicated a p-value less than or equal to 0.001.

Prior to any two-sample t-tests for mean differences, we evaluated the ratio of sample variances using Fischer’s F-test. If the F-statistic was less than or equal to three, sample variances were considered equal, and we used a Student’s t-test. Otherwise, we assumed sample variances were unequal, and we used Welch’s t-test. ANOVAs (two-way, one-way) were performed with post-hoc pairwise comparisons of means using a Tukey-Kramer adjustment.

### Exclusion criteria

#### Combining rotation conditions

In practice, we tested both positive and negative rotation conditions. However, we combined the positive and negative rotations based on the lack of statistical support for separating the conditions in both subjects. We investigated the main and interaction effects of rotation sign and perturbation timepoint on behavioral performance.

A two-way ANOVA with factors of rotation condition (levels: -50°, +50°) and timepoint (levels: initial drop, initial recovery, maximum recovery) revealed no significant main effects for the sign (Monkey A: F_1,90_ = 0.41, p=5.2e-01; Monkey B: F_1,75_ = 1.93, p=1.7e-01; main effect of sign). As expected, the main effects were limited to the perturbation timepoints (Monkey A: F_2,90_ = 53.99, p=3.9e-16; Monkey B: F_2,75_ = 52.63, p=5.2e-15, main effects of timepoint) with no significant interaction effect for Monkey A (F_2,90_ = 0.01, p=9.9e-01) and a trending interaction effect for Monkey B (F_2,75_ = 2.99, p=5.6e-02).

A two-way ANOVA with factors of rotation condition (levels: -90°, +90°) and timepoint (levels: initial drop, initial recovery, maximum recovery) revealed a trending main effects for the sign for Monkey A (F_1,87_ = 3.05, p=8.4e-02; main effect of sign) and a non-significant main effects for the sign for Monkey B (F_1,84_ = 0.97, p=3.3e-01; main effect of sign). As expected, the main effects were limited to the perturbation timepoints (Monkey A: F_2,87_ = 59.17, p=6.0e-17; Monkey B: F_2,84_ = 56.17, p=3.3e-16, main effects of timepoint) with no significant interaction effect (Monkey A: F_2,87_ = 0.68, p=5.1e-01; Monkey B: F_2,84_ = 0.80, p=4.5e-01).

#### Sessions and trials

The minimum inclusion criterion for a session was that the subject had to complete the first (baseline) and second (perturbation) blocks. Each of these blocks contains 384 trials. The baseline block included “error clamp” (EC) trials in which the cursor error feedback was projected (“clamped”) onto a straight-line path to the target. These trials, which occurred once every eight trials, were excluded from analyses in this paper. On some days, subjects would enter a third block (perturbation 2) in which error clamp trials were reinstated or a fourth block (washout) which reinstated the baseline decoder. The third and fourth blocks were not included for analyses; therefore, the following number of trials (on average) were not analyzed in this work: Monkey A completed, on average, 241.44 ± 136.42 trials past the first perturbation block for the easy rotation condition and 130.29 ± 132.26 trials past the first perturbation block for the hard rotation condition. Monkey B completed, on average, 235.87 ± 109.69 trials past the first perturbation block for the easy rotation condition and 237.27 ± 104.08 trials past the first perturbation block for the hard rotation condition.

Subjects were allowed to work until fluid satiation. Because both subjects completed trials beyond the baseline and perturbation blocks analyzed here, this suggests that satiation did not occur within the first two blocks. Therefore, they were motivated to work across the first two blocks.

#### Decoder units and factor analysis

To ensure that all decoder units were involved in the behavioral output, we excluded any units in which no spiking occurred in either the baseline or perturbation block. No units were excluded for Monkey A. For Monkey B, this excluded 2 units.

The average number of decoder units used in each rotation condition for neural analyses (Monkey A: *m*_*easy*_ = 36.28±7.28, *m*_*hard*_ = 37.65±6.89; Monkey B: *m*_*easy*_ = 97.56±13.44, *m*_*hard*_ = 91.90±26.59; mean ± standard deviation) did not significantly differ within subject (Monkey A: t_61_=-0.75, p=4.6e-01; Monkey B: t_55_=1.01, p=3.2e-01; two-sample, two-sided t-test, Monkey B: Welch’s alternative).

## Acknowledgements

H.M.S. was funded through the Department of Defense National Defense Science and Engineering Graduate Fellowship program. The authors would like to thank the Biomedical Imaging Center at the University of Texas at Austin for assistance with their MRI scanner.

